# Ketosis regulates K^+^ ion channels, strengthening brain-wide signaling disrupted by age

**DOI:** 10.1101/2023.05.10.540257

**Authors:** Helena van Nieuwenhuizen, Anthony G. Chesebro, Claire Polizu, Kieran Clarke, Helmut H. Strey, Corey Weistuch, Lilianne R. Mujica-Parodi

## Abstract

Aging is associated with impaired signaling between brain regions when measured using resting-state fMRI. This age-related destabilization and desynchronization of brain networks reverses itself when the brain switches from metabolizing glucose to ketones. Here, we probe the mechanistic basis for these effects. First, we established their neuronal basis using two datasets acquired from resting-state EEG (*Lifespan:* standard diet, 20-80 years, N = 201; *Metabolic:* individually weight-dosed and calorically-matched glucose and ketone ester challenge, *μ*_*age*_ = 26.9 ± 11.2 years, N = 36). Then, using the multi-scale Larter-Breakspear neural mass model, we identified the unique set of mechanistic parameters consistent with our clinical data. Together, our results implicate potassium (K^+^) gradient dysregulation as a mechanism for age-related neural desynchronization and its reversal with ketosis, the latter finding of which is consistent with direct measurement of ion channels.

## Introduction

Endogenous ketone bodies, including D-*β*-hydroxybutyrate (D-*β*HB), are produced by the liver and can be utilized by cells as fuel when glucose is not readily available [1]. Accumulating evidence suggests that ketone metabolism may confer important neurological benefits [2],[3],[4],[5]. D-*β*HB has been found to increase ATP production and cardiac efficiency from 10% to 24% when added to a perfusion of glucose [6]. Moreover, ketone uptake bypasses GLUT4, and thus can be metabolized as fuel even after neurons become insulin resistant [7].

The ability of insulin-resistant neurons to metabolize ketones is especially important for brain aging because Type 2 diabetes mellitus and its associated decrease in GLUT4-dependent neuronal glucose utilization are thought to be major contributors to age-related brain hypometabolism [8, 9] and cognitive decline [10, 11]. Indeed, even after aging brains show impaired glucose metabolism, they continue to be able to metabolize ketone bodies [12]. Thus, providing hypometabolic neurons with ketone bodies as an alternative fuel source may halt neurodegeneration [13]. We have previously shown, using resting-state fMRI, that the stability of functional communication across brain regions over time (*brain network stability*) changes in response to both longer-term dietary conditions (*fasting, glycolytic*, and *ketogenic*), as well as acute oral administration of individually weight-dosed D-*β*HB versus calorically matched glucose. Specifically, under glycolytic conditions, brain aging is associated with the destabilization of brain networks, which–even in healthy young adults–stabilize under ketosis, regardless of whether ketosis was induced by diet or administration of exogeous ketones [14]. We further measured a collective measure of brain activity (*brain network synchrony*) to decrease with age, which likewise increased with ketosis [15]. However, both aging [16] and ketosis [17] affect cerebral blood flow, and therefore also fMRI’s assumptions regarding neuro-vascular coupling [18]. As such, it is unclear whether fMRI-derived network effects result from the modality’s neuronal versus hemodynamic influences.

Here, we significantly extend our prior findings to focus on mechanism, proceeding in two stages. At the first stage, we used EEG, a measure of neuronal but not hemodynamic activity, to determine whether age and metabolism-related changes in brain network stability and synchrony do, in fact, reflect neuronal influences. To test the impact of aging, we used a *Lifespan* resting-state EEG dataset (ages 20-80 years, N = 201) to measure brain network stability and synchrony. Holding age constant, we then isolated the impact of metabolism alone. In an independent *Metabolic* resting-state EEG experiment (*μ*_*age*_ = 26.9 ±11.2 years, N = 36) (Fig 1A), we orally administered individually weight-dosed D-*β*HB versus calorically matched glucose, measuring their impact on these same network variables (Figs. 1B, 1C). After establishing that age and metabolic modulation of brain networks were neuronal effects, we used the Larter-Breakspear multi-scale neural mass model [19] to identify the unique set of mechanisms at the neuronal micro-scale consistent with these measured differences in EEG-derived neural synchrony at the macro-scale Fig. 1D).

**Figure 1.**
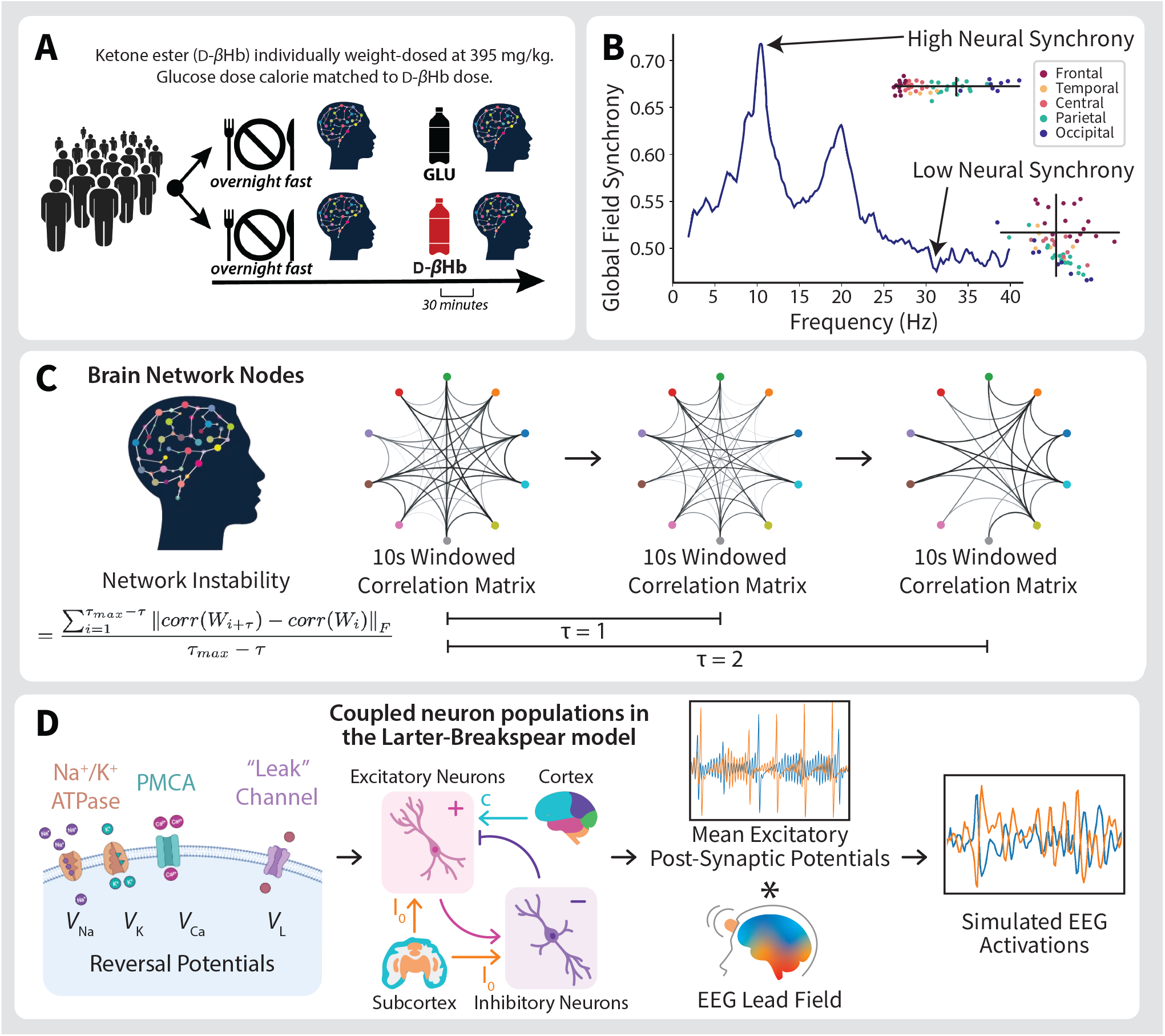
Schematic of experimental design and methods. (A) *Design of within-subject, time-locked targeted metabolic resting-state EEG (rsEEG) experiment*. To confirm the neuronal origins of increased brain network stability and synchrony, N = 36 young (*μ*_*age*_ = 26.9 ± 11.2 years), healthy participants underwent four rsEEG scans separated over two days. Following an overnight fast, participants were scanned at baseline and again 30 minutes after consuming a weight-dosed (395 mg/kg) ketone ester or calorie-matched glucose bolus. The rsEEG scans were then repeated using the opposite (ketotic or glycolytic) condition on the second day. (B) *An example Global Field Synchronization (GFS) spectrum computed using human rsEEG*. The real and imaginary components of Fourier-decomposed rsEEG time-series are plotted on the complex plane for each frequency value. The spread of these points is quantified using principal component analysis. The more the signals are in phase or anti-phase, the greater the difference in magnitude between the first and second principal components of the scatter plot cloud, and the greater the synchrony value (can range from 0 to 1). The scatter points of individual electrodes are color-coded by location on the scalp for illustrative purposes. (C) *Schematic characterization of brain network instability*. To calculate brain network instability, non-overlapping sliding window correlations are calculated over the entire rsEEG time-series, with strong correlations defining networks. The stability of the networks is then defined as the degree to which these networks persist over time (in units of *τ*). (D) *Schematic of the Larter-Breakspear neural mass model*. Microscale parameters (listed in Table 1), along with intra-/inter-region coupling (*c*) and subcortical excitatory inputs (*I*_0_) govern the dynamics of the model output: simulated excitatory post-synaptic potentials (EPSPs). The mean EPSPs are multiplied with an EEG lead field to generate simulated EEG time-series, which are used to determine the effects of model parameter variation on synchrony.

## Results

### Age modulates neural signaling

Using the Leipzig LEMON rsEEG dataset, a comparison of Global Field Synchronization (GFS, see Methods) between the younger (aged 20 to 40 years, N = 138) and older (aged 55 to 80 years, N = 63) cohorts within classically defined frequency bands (*δ* (1 - 3.5 Hz), *θ* (4 - 7.5 Hz), *α* (8 - 13 Hz), *β* (14 - 30 Hz), low *γ* (30 - 40 Hz), and broad (1 - 40 Hz), [20]) showed age to decrease neural synchrony in the broad, *θ, α*, and low *γ* bands (Fig. 2A). Likewise, older brains showed destabilization of their networks (Fig. 2C).

**Table 1.** Parameter values used in the Larter-Breakspear neural mass model. The “Default Value” column lists the default parameter values used in the Larter-Breakspear model, following Endo et al. [45] and Chesebro et al. [24]. If the parameter was varied in order to examine the effects on synchrony, the range of parameter values tested is listed in the “Variation Range” column. Likewise, the step size used to sample between lowest and highest possible varied parameter value is listed in the “Step Size” column.

| Parameter | Description | Default Value | Variation Range | Step Size |
| --- | --- | --- | --- | --- |
| **Nernst potentials** |  |  |  |  |
| $V_{Na}$ | $Na^+$ reversal potential | 0.53 | [0.42, 0.54] | 0.005 |
| $V_K$ | $K^+$ reversal potential | -0.7 | [-0.75, -0.6575] | 0.0025 |
| $V_{Ca}$ | $Ca^{2+}$ reversal potential | 1.0 | [0.95, 1.01] | 0.0025 |
| $V_L$ | Leak channels reversal potential | -0.5 | | |
| **Channel conductances** |  |  |  |  |
| $g_{Na}$ | $Na^+$ conductance | 6.70 | [6.6, 6.8] | 0.01 |
| $g_K$ | $K^+$ conductance | 2.00 | [1.95, 2.05] | 0.0025 |
| $g_{Ca}$ | $Ca^{2+}$ conductance | 1.00 | [0.95, 1.01] | 0.0025 |
| $g_L$ | Leak channels conductance | -0.50 | | |
| **Channel voltage thresholds** |  |  |  |  |
| $T_{Na}$ | $Na^+$ channel threshold | 0.30 | | |
| $T_K$ | $K^+$ channel threshold | 0.00 | | |
| $T_{Ca}$ | $Ca^{2+}$ channel threshold | -0.01 | | |
| **Channel voltage threshold variances** |  |  |  |  |
| $\delta_{Na}$ | $Na^+$ channel threshold variance | 0.15 | | |
| $\delta_K$ | $K^+$ channel threshold variance | 0.30 | | |
| $\delta_{Ca}$ | $Ca^{2+}$ channel threshold variance | 0.15 | | |
| **Excitatory & inhibitory population parameters** |  |  |  |  |
| $V_T$ | Excitatory neuron threshold voltage | 0.00 | | |
| $Z_T$ | Inhibitory neuron threshold voltage | 0.00 | | |
| $\delta_{V,Z}$ | Variance of excitatory/inhibitory thresholds | 0.66 | | |
| $Q_{V_{max}}$ | Excitatory population maximum firing rate | 1.00 | | |
| $Q_{Z_{max}}$ | Inhibitory population maximum firing rate | 1.00 | | |
| $a_{ee}$ | Excitatory → excitatory synaptic strength | 0.36 | [0.33, 0.39] | 0.005 |
| $a_{ei}$ | Excitatory → inhibitory synaptic strength | 2.00 | [1.95, 2.05] | 0.005 |
| $a_{ie}$ | Inhibitory → excitatory synaptic strength | 2.00 | | |
| $a_{ne}$ | Non-specific → excitatory synaptic strength | 1.00 | | |
| $a_{ni}$ | Non-specific → inhibitory synaptic strength | 0.40 | | |
| **Other parameters** |  |  |  |  |
| $I_0$ | Subcortical excitatory input | 0.30 | | |
| $b$ | Time scaling factor | 0.10 | | |
| $\phi$ | Temperature scaling factor | 0.70 | | |
| $\tau_K$ | $K^+$ relaxation time | 1.00 | | |
| $r_{NMDA}$ | NMDA/AMPA receptor ratio | 0.25 | | |
| $c$ | ROI-to-ROI coupling constant | 0.35 | | |

**Figure 2.**
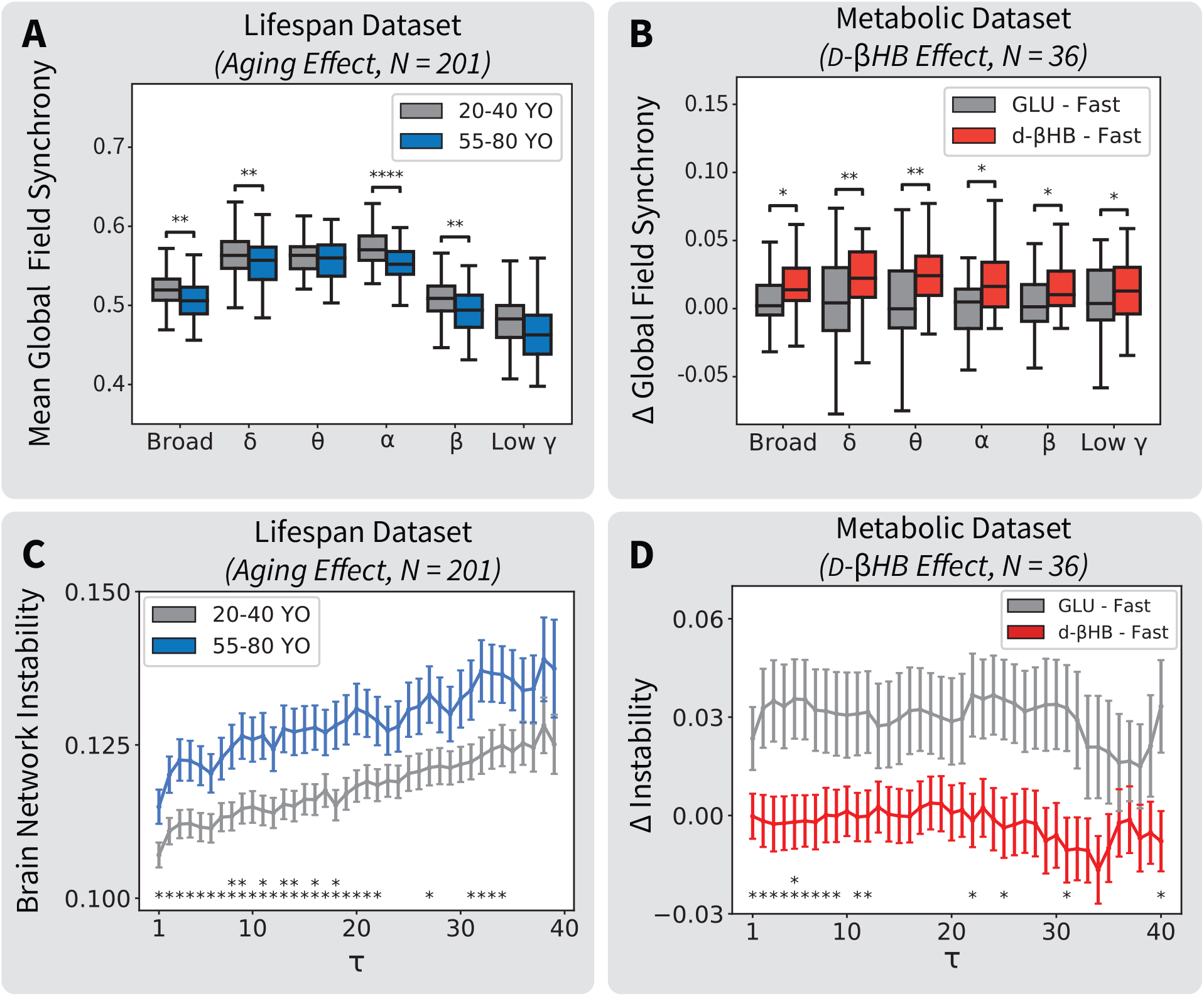
Global Field Synchronization, a global measure of neural synchrony, decreases with age and increases under ketosis. Brain network instability, a measure of dynamic functional connectivity, increases with age, and decreases under ketosis. (A) Mean synchrony is lower in the older (N = 63, 55 to 80 years) cohort than the younger (N = 138, 20 to 40 years) cohort in the broad (t = −3.260, p = 0.001), *δ* (t = −3.001, p = 0.003), *α* (t = −5.120, p *<* 0.001), and *β* (t = −2.965, p = 0.003) bands of Leipzig’s LEMON EO, rsEEG scans. (B) Baseline-corrected synchrony is greater following acute administration of D-*β*HB when compared to acute administration of glucose in all classically-defined frequency bands of EO, rsEEG scans: broad (t = 2.659, p = 0.012), *δ* (t = 2.763, p = 0.009), *θ* (t = 3.184, p = 0.003, *α* (t = 2.693, p = 0.010), *β* (t = 2.331, p = 0.026), and low *γ* (t = 2.235, p = 0.032), measured using a cohort of young, healthy individuals (N = 36, *μ*_*age*_ = 26.9 *±* 11.2 years). (C) Brain network instability (window size = 10 s) is significantly greater in the older cohort than the younger cohort in the broad band of Leipzig’s LEMON EC, rsEEG scans for 27 out of 39 possible values of *τ*, with p-values ranging from p = 0.005 to p = 0.045 and t-values ranging from t = 2.023 to t = 2.846. When comparing across bandwidths, brain network instability (window size = 10 s, *τ* = 1) of the older cohort is significantly greater in the broad (t = 2.278, p = 0.024), *δ* (t = 3.581, p *<* 0.001), *θ* (t = 2.323, p = 0.021), and low *γ* (t = 3.513, p *<* 0.001) bands. (D) Baseline-corrected brain network instability (window size = 10 s) is significantly greater after acute administration of glucose when compared to acute administration of ketones in the broad (1-40 Hz) band of EC, resting-state EEG scans performed on a cohort of young, healthy individuals (N = 36, *μ*_*age*_ = 26.9 ±11.2 years) for 15 out of 40 possible values of *τ*, with p-values ranging from p = 0.009 to p = 0.047 and t-values ranging from t = 2.055 to t = 2.744. The stabilization of brain networks under ketosis was not dominated by a particular frequency band. Baseline correction for both metrics was performed by subtracting the pre-bolus (fasting) condition value from the post-bolus (either glucose or D-*β*HB) condition value within each respective metric.

### Metabolism modulates neural signaling

Changes in GFS values (Δ GFS) for the metabolic study cohort (N = 36, *μ*_*age*_ = 26.9 ±11.2 years) were calculated by subtracting the average fasting condition GFS from the average experimental (glycolytic or ketotic) condition GFS within each band. Comparing the pre- and post-bolus synchrony values showed decreased synchrony after administration of the glucose bolus and increased synchrony after administration of the ketone bolus for all frequency bands in the rsEEG eyes-open (EO) condition (Fig. 2B). Following glucose challenge, brain networks destabilized; in contrast, following ketone challenge, brain networks stabilized. (Fig. 2D) in rsEEG. Both results were observed in the broad (1-40 Hz) frequency band of eyes-closed (EC) rsEEG.

### Identifying candidate mechanisms using multi-scale modeling

We next sought to identify the mechanistic basis for age and metabolism-related changes observed in rsEEG GFS. To do so, we used a bottom-up approach by varying microscale parameters in the Larter-Breakspear neural mass model to generate 2,082 simulated rsEEG signals, from which we computed GFS and instability. We observed high sensitivity of GFS to changes in all (Ca^2+^, K^+^, and Na^+^) Nernst potentials, Ca^2+^ and K^+^ channel conductances, and excitatory → inhibitory synaptic strength (Fig. 3B). Only variation of the K^+^ Nernst potential within this model explained both the magnitude of the changes seen with older age (2.5% decrease) and following D-*β*HB consumption (5.6% increase) in broadband GFS (Fig. 3C). Sensitivity of brain network instability to the parameter changes was binary: either it was not sensitive to changes in parameters, or network instability changed in non-consistent ways and with large magnitudes that did not reflect the smaller magnitudes seen in human data (Fig. S1). Furthermore, brain network instability was found to be uncorrelated with GFS within the broad (1-40 Hz) frequency band of Leipzig’s LEMON rsEEG dataset (p = 0.090, *r*^2^= 0.014, N = 201).

**Figure 3.**
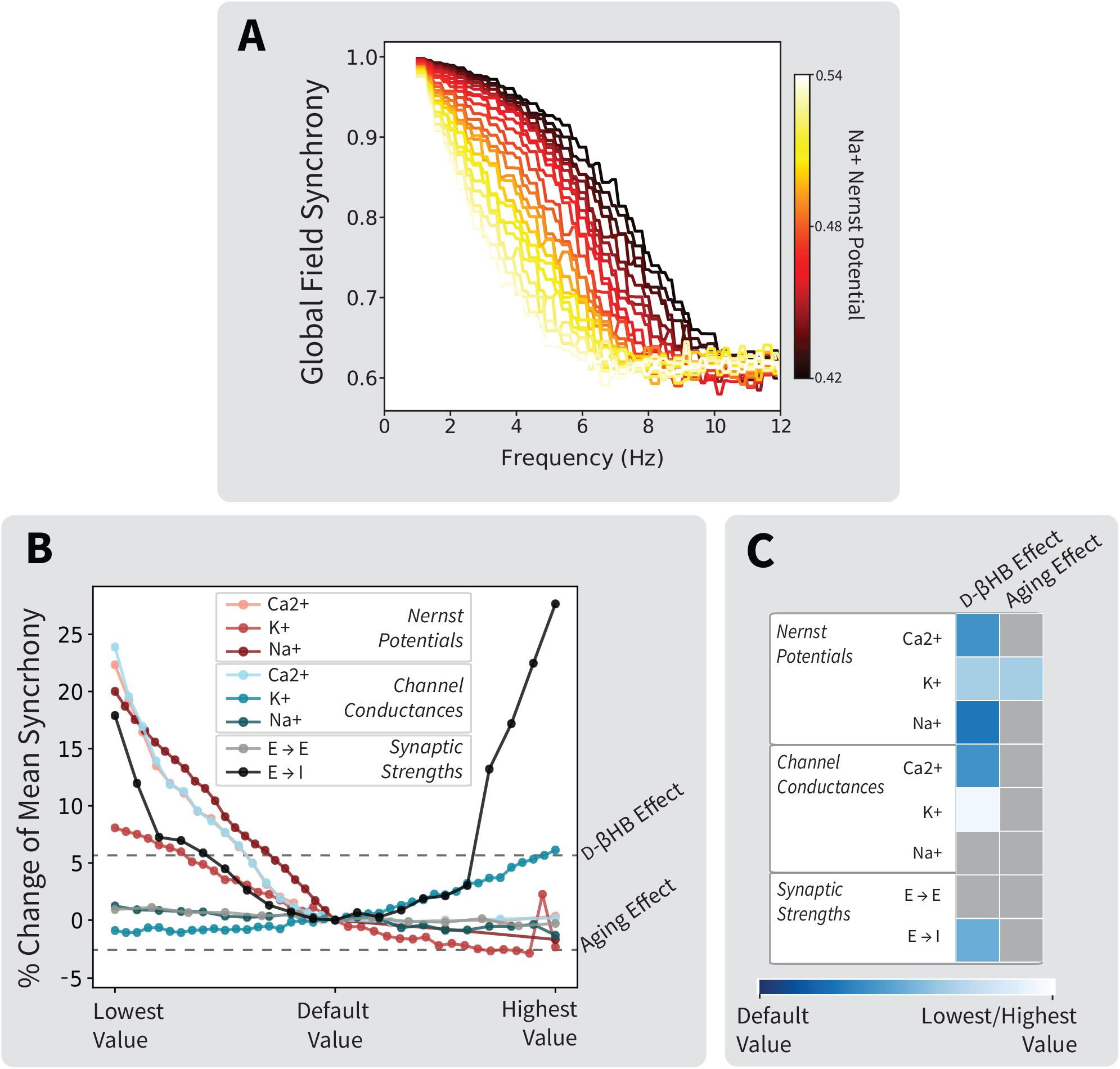
Variation of the K^+^ Nernst potential within the Larter-Breakspear neural mass model uniquely predicts changes in Global Field Sychrony (GFS) seen in aging and following acute administration of D-*β*HB. (A) An example of the GFS spectra for simulated EEG data generated using the Larter-Breakspear neural mass model. The color indicates the value of the Na^+^ Nernst potential used in the simulation. (B) Individually varying parameters within the model above and below their default values leads to changes in GFS. Parameters varied are color-coded by class (Nernst potentials, channel conductances, and excitatory → excitatory/inhibitory synaptic strengths). (C) Heatmap of the relative parameter change needed to replicate the magnitude of changes seen in broadband GFS (Figs. 2A, 2B).

## Discussion

It has been hypothesized that brain aging results from an “energy crisis” in the brain, in which decreased glucose oxidation capability leads to constrained ATP availability for neurons [21],[22], disrupting neuronal signaling, and thus dysregulating the neural circuits that underlie cognitive processing. Using the Larter-Breakspear model, parameter sensitivity analyses identified modulation of the K^+^ Nernst potential as the only mechanism consistent with the macroscale synchrony age and metabolism-related changes we measured in rsEEG. The fact that our results independently match results obtained by *in vitro* electrophysiology, confirming that ketosis modulates neuron K^+^ regulation [23], increases confidence in the validity our multi-scale approach.

Dysfunction of Na^+^/K^+^-ATPase causes depolarization of the membrane potential, and thus desynchronization between brain regions [24]. Ion pumps are a sink of metabolic resources in the brain, and thus Na^+^/K^+^-ATP-ase dysregulation is also expected under metabolic constraints, which would further exacerbate ion gradient dysregulation [25]. Changes in potassium reversal potential have been implicated in a number of different age-related processes [26],[27]. The depletion of the potassium gradient can stem from damage to potassium channels by reactive oxygen species [26] or from calcium signaling dysregulation [28],[29], or by a combination of accumulated insults, leading to impairment.

The Larter-Breakspear model provides a biophysically-detailed simulation of ion dynamics while allowing for a whole-brain simulation by abstracting away population details. While this derivation preserves local ion dynamics [30], it does not fully capture emergent properties of neural populations (e.g., the model preserves cortical wave dynamics [31], but not power spectra). We found that GFS was sensitive to changes in the biophysical model parameters(Fig. 3B), reflecting changes that were on the same scale of magnitude as the changes in GFS seen in the human (non-simulated) data. However, the absolute magnitude of the spectra produced using our modeling approach did not match that of spectra produced from our human rsEEG data. Furthermore, brain network instability was either not sensitive at all to changes in parameters, or for those parameters that did alter network instability, changed it in non-consistent ways and with large magnitudes of change that did not accurately reflect the smaller magnitudes seen in human data (Fig. S1). These limitations may be due to the inability of the Larter-Breakspear neural mass model to capture certain emergent properties of neural populations, and have been partially addressed in next-generation neural mass models [32]. Brain network instability and neural synchrony were found to be uncorrelated, implying the two metrics quantify distinct neural features, of which the features of the latter are better captured by the Larter-Breakspear neural mass model. Next-generation neural mass models may be able to shed light on the similarities and differences between various metrics used in the neuroimaging community. As of yet, however, these models do not incorporate the same biophysical detail as the Larter-Breakspear model, and fusing these approaches is a topic of current interest for the field of computational neuroscience.

Prior literature has established ion gradient regulation to be the most significant energy sink within the brain [25], [33]. This is consistent with our results linking emergent whole-brain network effects to ion gradient regulation. Our further isolation of the consistent candidate mechanisms to K^+^ Nernst potentials is also in agreement with *in vitro* experimental evidence that ketosis directly impacts neuronal K^+^ regulation [23]. Together, results across scales suggest future directions in directly testing the hypothesis that age-based hypometabolism is associated with neuronal K^+^ gradient dysregulation, reversed through ketosis. If correct, it would demonstrate that generative models constrained by clinical neuroimaging data can bridge the gap between molecularly-targeted treatments and patient outcomes.

## Acknowledgments

The research described in this paper was funded by the White House Brain Research Through Advancing Innovative Technologies (BRAIN) Initiative (grants NSFNCS-FR 1926781 to LRMP) and the Baszucki Brain Research Fund, United States (LRMP). CW acknowledges support from the Marie-Josée Kravis Fellowship in Quantitative Biology. AGC was also supported by the NIHGM MSTP Training Award, United States T32-GM008444.

## Author Contributions

LRMP, CW, and KC designed research. HvN and CP performed research and collected data. HvN analyzed data. HvN, AC, and HHS performed computational modeling. KC contributed reagents. HvN, LRMP, and AC wrote the paper. HvN, AGC, KC, HHS, CW, and LRMP edited the paper.

## Competing Interest Statement

The intellectual property covering the uses of ketone bodies and ketone esters is owned by BTG Plc., Oxford University Innovation Ltd., and the NIH. KC, as inventor, will receive a share of the royalties under the terms prescribed by each institution. KC is a director of TΔS Ltd., a company spun out of the University of Oxford to develop products based on the science of ketone bodies in human nutrition.

## Data and Code Availability

Data and code are available at our website: https://www.lcneuro.org/data/eeg. Leipzig LEMON data is publicly available at http://fcon_1000.projects.nitrc.org/indi/retro/MPI_LEMON.html.

## Methods

### Bolus Study Experimental Design

To determine whether stabilization of brain networks as modulated by fuel source seen in rsfMRI [14],[15] reflect a neuronal, rather than only hemodynamic, response we conducted resting-state EEG scans on a cohort of healthy adults (*N* = 36, *μ*_*age*_ = 26.9 ±11.2 years, 18 female) who were tested under both a ketotic (ketone burning) and glycolytic (glucose burning) condition. Inclusion/exclusion criteria was screened using a survey completed prior to enrollment in the study. Potential participants were excluded for any of the following reasons: chronic usage of alcohol (7 drinks/week for women, 14 drinks/week for men), recreational drug use, use of psychotropic medication within the past 30 days, use of medications that affect glucose and/or insulin utilization, a history of kidney disease, heart attack, stroke, myxedema, epilepsy, dementia, or other neurological disorders, difficulty when swallowing, current pregnancy or breastfeeding, or inability to provide informed consent. Scanning occurred at Stony Brook University’s Health Science Center. The study was approved by the Institutional Review Board of Stony Brook University (IRB2021-00018) and all participants provided informed consent.

All subjects were tested twice (1-14 days apart), both times following an overnight fast (subjects were instructed to eat no food for at least 8 hours prior to testing, but were allowed unrestricted water). Following a baseline resting-state EEG scan, subjects drank either of two fuel sources. In the ketotic session subjects drank a ketone sports drink, deltaG® Sports Supplement (TdeltaS Ltd, Thame, UK), dosed at 395 mg/kg. During the glycolytic session, the same subjects drank a bolus of glucose (Glucose Tolerance Test Beverages, Fisher Scientific, Inc.; Hampton NH) calorie-matched to the ketone bolus. The order of the bolus (whether a subject received a glucose bolus first or a ketone bolus first) was pseudo-randomized, with approximately half of all female subjects (N = 9) and exactly half of all male subjects (N = 8) drinking the glucose bolus during the first scanning day. The resting-state EEG scans were then repeated 30 minutes after the administration of the bolus, as prior experiments using magnetic resonance spectroscopy (MRS) showed peak glucose and ketone concentration in the brain approximately 30 minutes after consumption of the bolus. Blood glucose and ketone levels were measured three times throughout the duration of the experiment - at baseline, 10 minutes following the bolus, and 62 minutes following the bolus using a Precision Xtra Blood Glucose & Ketone Monitoring System (Abbott Laboratories). Our data analyses quantify network reorganization and neural phase synchrony changes in response to changing energy constraints (i.e., cognitive demand, fuel).

During the resting-state portion of the EEG scan, subjects underwent a total of 16 blocks, each lasting 60 seconds, with 8 EO blocks and 8 EC blocks. The blocks were interleaved. During the EO blocks, subjects fixated on a white orienting cross on a black background. Prior to the resting-state scan subjects were instructed not to blink in order to minimize ocular artifacts and to keep motion to a minimum. The resting-state stimulus (a white cross in the center of a black background) was presented on a computer screen placed in front of the seated subject using PsychoPy v3.0 [34]. All data were collected in a shielded, dark, soundproofed Faraday cage using the ActiveTwo Biosemi™ electrode system from 65 (64 scalp, 1 ocular) electrodes arranged according to the International standard 10-20 system [35] at a sampling frequency of 4096 Hz. The ocular (VOEG) electrode was placed below the left eye. Our experimental and pre-processing designs, especially those of the resting-state EEG scans, were modeled after the paradigm used by Leipzig’s LEMON rsEEG dataset group [36] in order to be able to minimize confounding factors when directly comparing results between our experiment and this large-scale neuroimaging dataset.

### Resting-state EEG Pre-Processing

#### Metabolic Dataset

The EEG pre-processing was performed using EEGLab (version 2020.0) [37]. Full resting state data were downsampled from 4096 Hz to 512 Hz and bandpass filtered between 0.1 and 40 Hz using a Hamming-windowed FIR filter. The data were then separated into the eyes open (EO) and eyes closed (EC) conditions. These two conditions were pre-processed separately from this point on due to the differences in ocular artifacts in each condition, however, the pre-processing steps performed were identical. Bad channels were removed and noisy portions of data were identified and removed using EEGLab’s Artifact Subspace Reconstruction (ASR) algorithm. Independent component analysis (ICA) was performed on the data using the infomax algorithm in EEGLab (runica), and non-neural components of the time-series were identified and removed using ICLabel [38]. The reference was then set to average. The data were separated into frequency bands and time-series extracted using MNE-Python (version 0.21.1) [39].

#### Leipzig LEMON Dataset

Lifespan rsEEG data from the Leipzig LEMON dataset were obtained in already pre-processed form. Raw data were downsampled from 2500 Hz to 250 Hz and bandpass filtered between 1 and 45 Hz using an 8th order Butterworth filter. The data were separated into EO and EC conditions for subsequent pre-processing. Outlier channels were rejected and data intervals containing high peak-to-peak fluctuations or high-frequency noise were identified and removed by visual inspection. Data dimensionality was reduced using principal component analysis prior to the use of independent component analysis (ICA) to identify and remove components reflecting eye-or heartbeat-related artifacts. Further pre-processing details may be found in the dataset documentation [36].

### Global Field Synchronization

Global Field Synchronization was first introduced by König et al. [40] to estimate differences in functional connectivity of brain processes in EEG frequency bands between a population of neuroleptic-naïve schizophrenic patients and healthy controls. In contrast to other measures of phase synchrony such as phase-locking value, which can only measure synchrony between two time-series, GFS is a global measure of neural phase synchrony that does not rely on the a priori selection of brain regions to be studied. When applied to EEG data, GFS has the added benefit of being reference-independent and more easily interpretable without the use of source models [41]. Further research using the measure has found changes in GFS in those with Alzheimer’s disease and mild cognitive impairment [42], during REM sleep [43], and during general anaesthesia [44].

In order to calculate GFS values for EEG, the data were first pre-processed. Following pre-processing, sensor-level EEG time-series were divided into non-overlapping, consecutive 2 second epochs. Each epoch was frequency transformed using a fast Fourier transform (FFT, Tukey window, taper size *α* = 0.2), which yields the real and complex component of the signal of each electrode for each frequency value (1 - 40 Hz, step size = 0.1 Hz). These components are then plotted as vectors on the complex plane, with the magnitude of the vector representing the power of the signal at that frequency, the angle of the vector as measured from the real axis representing the phase, and the vector origin representing the reference of all EEG signals. Subsequently, a scatter plot of vector endpoints in the complex plane is generated for each frequency value, a representative of which can be seen in figure 1B. The more these endpoints approximate a straight line, the more the signals are in phase or anti-phase. The more scattered the endpoints, the less the signals are in phase or anti-phase. In order to quantify the shape of the cloud formed by the vector endpoints, the points are entered into a two-dimensional principal component analysis, as principal components are orthogonal by definition. The resulting GFS value per epoch for a particular frequency *f* is then determined by calculating the ratio of the eigenvalues (*λ*_1_(*f*) and *λ*_2_(*f*)) of these two principal components, as expressed in equation 1.

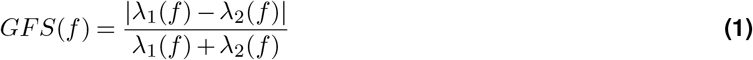

Finally, the overall GFS value for a particular frequency is obtained by taking the mean of the GFS values of all consecutive epochs for that frequency, creating a spectrum as shown in figure 1B. In order to examine differences in GFS across age (using the Leipzig LEMON rsEEG dataset) and condition (ketotic vs. glycolytic), overall GFS values were categorized into their corresponding, classically-accepted frequency bands [20], with endpoints adjusted to accommodate the filtering performed in pre-processing: *δ* (1 - 3.5 Hz), *θ* (4 - 7.5 Hz), *α* (8 - 13 Hz), *β* (14 - 30 Hz), low *γ* (30 - 40 Hz), and broad (1 - 40 Hz).

### Modeling Effects of Microscale Parameter Changes on Global Field Synchronization

We used the Larter-Breakspear neural mass model [19] to test the effect of ion gradient dynamics and excitatory-inhibitory subpopulation coupling on GFS. The Larter-Breakspear model has been previously validated to capture properties of fMRI and EEG resting-state activity [45] while incorporating both local ion dynamics and interregional connectivity. Following the mathematics of prior work [45], we built a 78-region whole-brain simulation using Neuroblox, a Julia library optimized for high-performance computing of dynamical brain circuit models (https://github.com/Neuroblox/Neuroblox.jl). Within the model, we varied ion (Na^+^, K^+^, Ca^2+^) gradients and conductances, intraregional connectivity (excitatory-excitatory/inhibitory), and interregional connectivity across a range of biophysically plausible values [24],[31]. Further details, including equations, can be found in the methods. The whole-brain simulation for each parameter set was repeated with 10 different sets of initial conditions in order to sample across the distribution of simulation outcomes. The simulated membrane potentials of excitatory neurons were averaged within each region to generate 78 neural mass signals, which were transformed into EEG signals through multiplication with an average lead field (63 EEG channels × 78 model ROIs) generated by Endo et. al [45]. Each 10-minute simulation was sampled at a rate of 1000 Hz and took ∼2 minutes to generate. The first 1 minute of each simulated EEG signal was discarded prior to calculating GFS in order to allow the simulation to equilibrate.

### Brain Network Instability

Brain network instability (described for use with fMRI data in [14]) is a measure used to describe the persistence of brain networks over time, and can be considered a measure of dynamic functional connectivity [46]. In order to calculate brain network instability values for EEG, the data were first pre-processed as described above. Following pre-processing, sensor-level EEG time-series were divided into non-overlapping, consecutive epochs, or windows, with a 10 s window chosen for the results shown in Figs. 2C and 2D. Within each window *W*_*i*_ an all-to-all, signed correlation matrix between all time-series is calculated, meaning these resulting matrices have a size of *n × n*, with *n* being the number of clean channels remaining after pre-processing. Distance (in units of window size) between window pairs is decided by a value of *τ* chosen for the instability calculation. Instability is then calculated for each possible value of *τ* by taking the Frobenius, or L2, norm of the difference in correlation matrices of window pairs, and then taking the average of all norms. In the form of an equation, instability is written as

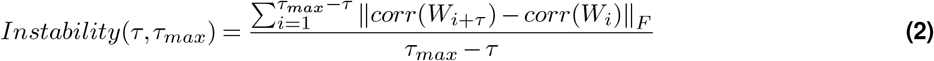

where *τ*_*max*_ is the number of windows available, defined as the length of the time-series divided by the window size rounded down to the nearest multiple of the window size. For example, an EEG recording with a length of 55 s divided into window sizes of 10 s means *τ*_*max*_ = 5. Computing instability for *τ* = 2 for this example recording would yield

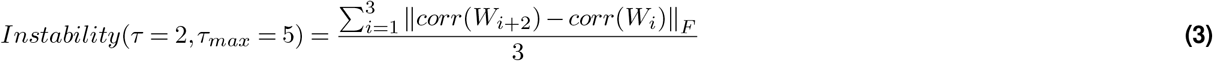

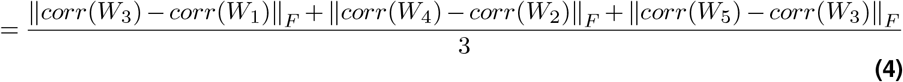

When calculating instability, the window sizes within which correlation matrices are calculated and eventually subtracted from one another set a natural limit to the timescale at which changes in global network connectivity can be seen. As the temporal resolution of EEG is notably greater than that of fMRI, the window sizes used in the calculation of instability for the majority of this work are smaller than the windows used for our previous work (10s as compared to 24s) as more time points are available for calculation. By varying the window sizes used to calculate network instability, we determined that brain networks destabilize as a function of age only for networks whose detectable correlation differences persist for ≥10 s (Fig. S2), and as such made the choice of window size to be 10 s for subsequent analyses. Furthermore, to ensure the increase in instability with age seen in the Leipzig LEMON dataset was not due to motion (as subject motion tends to increase with age [47]), we examined the relationship between the mean frame displacements (in mm) measured during the subjects’ corresponding rsfMRI scan and their brain network instability (*τ* = 1, window size = 10 s) within the broad (1-40 Hz) frequency band of the subjects’ EC, rsEEG scan. We found no correlation between these two variables.

### Larter-Breakspear Model: Single Neural Mass

Using the physiological and mathematical boundary conditions discussed in Chesebro et al. [24] to inform the model, the Larter-Breakspear equations are constructed as a system of three variables: mean excitatory membrane voltage *V* (*t*), mean inhibitory membrane voltage *Z*(*t*), and the proportion of potassium channels open at a given time *W* (*t*). Note that while *V* (*t*), *Z*(*t*), and *W* (*t*) are all time-dependent, we omit this dependence in the following equations for ease of reading. Given this understanding, the Larter-Breakspear model is defined as:

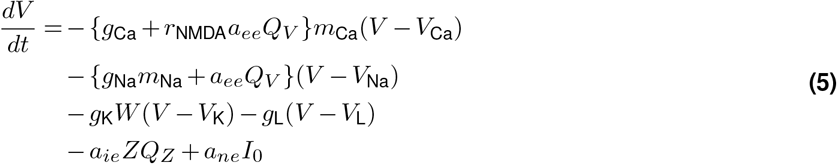

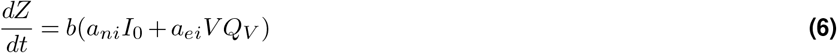

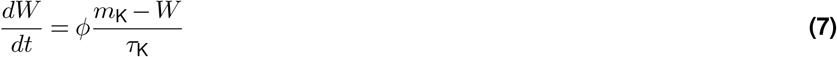

In these equations, *Q*_*V*_ and *Q*_*Z*_ are the mean firing rates for excitatory and inhibitory cell populations, respectively. These are computed as

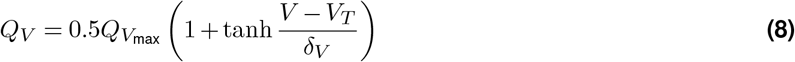

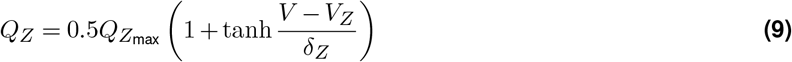

The individual ion channel gating functions (*m*_Na_, *m*_K_ and *m*_Ca_) take the form

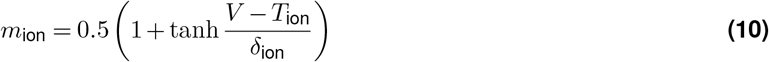

where *m*_ion_ is the fraction of voltage-dependent channels open at any given time. Default values and descriptions for all constants in these equations are given in Table 1. Note that parameter values are unit-less to scale to a reasonable modeling range (i.e., *V, Q ∈* (−1, 1) and *W ∈* (0, 1)), and the integration time step *dt* is in milliseconds.

We note that the three ions of interest are modeled in three different manners. Sodium serves as the dominant shape determinant of the neural mass spiking activity as it has the highest net positive conductance coupled to its ion channel gating function. Calcium serves as a secondary support of the spiking activity, providing some of the amplitude to the spiking activity. However, because the calcium gating function is also coupled to the excitatory population firing rate and the ratio of NMDA/AMPA receptors, calcium more importantly provides feedback to the neural mass firing rate. Finally, due to its more detailed modeling as a separate differential equation, potassium plays a unique role in determining the frequency of spiking activity (at larger reversal potentials) and the duration of the refractory period (at smaller reversal potentials). As a consequence of this extra modeling step, potassium also has a different ion gradient landscape than sodium and calcium.

While the excitatory (pyramidal) cell population *V* (*t*) is modeled using the ion dynamics described above, the inhibitory population *Z*(*t*) is a purely phenomenological model, receiving only the excitatory input via *a*_*ei*_ and a background current via *a*_*ni*_. Although this serves to model the relationship between inhibitory interneurons and the excitatory pyramidal cells (as in Larter et al. [30]), it does imply a caveat when interpreting the effects of altered ion gradients.

Since the model is a hybrid of a biophysically detailed excitatory neuron population and a phenomenological inhibitory population, claims regarding how closely this model resembles true biological neurons are necessarily constrained. However, the advantage of the Larter-Breakspear model is that it produces physiologically interesting dynamics (e.g., burst-spiking) that are more common in next generation neural mass models [48] than in traditional (e.g., Wilson-Cowan) oscillatory models.

### Larter-Breakspear Model: Coupled Neural Masses

Equations 5-7 describe a single neural mass comprised of two subnetworks. Coupling between pairs of neural masses (*i* and *j*) can also be achieved through connection terms:

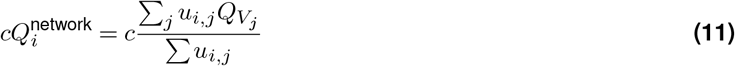

Here, *c* is the coupling constant, 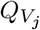 is the mean excitatory firing rate of region *j*, and *u*_*i,j*_ is the strength of connection between regions *i* and *j*. To ensure that overall input current is approximately constant, the balancing between interregional and self coupling takes the form of competitive agonism, where *c* is the weight of interregional coupling and (1 *-c*) is self-coupling. The associated multi-regional Larter-Breakspear equations are then given by:

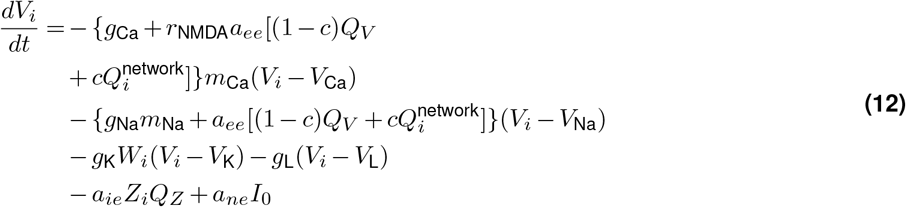

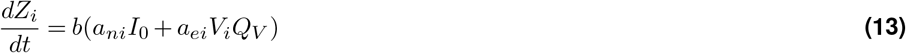

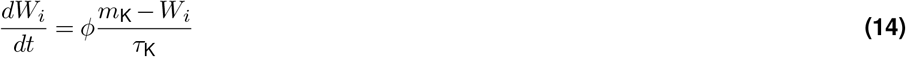

In this work, we use equations 12-14, varying the microscale parameters (Table 1) therein in order to observe the effects on macroscale neural synchrony. Following prior work [45], we computed a 78 region whole-brain model.

## Notes

http://www.lcneuro.org/data/eeg

## Bibliography

1. Krebs, H. Biochemical aspects of ketosis, 1960.

2. Barañano, K. W. and Hartman, A. L. The ketogenic diet: uses in epilepsy and other neurologic illnesses. Current treatment options in neurology, 10(6):410–419, 2008.

3. Henderson, S. T., Vogel, J. L., Barr, L. J., Garvin, F., Jones, J. J., and Costantini, L. C. Study of the ketogenic agent ac-1202 in mild to moderate alzheimer’s disease: a randomized, double-blind, placebo-controlled, multicenter trial. Nutrition & metabolism, 6(1):1–25, 2009.

4. VanItallie, T. B., Nonas, C., Di Rocco, A., Boyar, K., Hyams, K., and Heymsfield, S. Treatment of parkinson disease with diet-induced hyperketonemia: a feasibility study. Neurology, 64(4):728–730, 2005.

5. Maalouf, M., Sullivan, P. G., Davis, L., Kim, D. Y., and Rho, J. M. Ketones inhibit mitochondrial production of reactive oxygen species production following glutamate excitotoxicity by increasing nadh oxidation. Neuroscience, 145(1):256–264, 2007.

6. Sato, K., Kashiwaya, Y., Keon, C., Tsuchiya, N., King, M., Radda, G., Chance, B., Clarke, K., and Veech, R. L. Insulin, ketone bodies, and mitochondrial energy transduction. The FASEB Journal, 9(8):651–658, 1995.

7. Sapolsky, R. M. Glucocorticoid toxicity in the hippocampus: reversal by supplementation with brain fuels. Journal of Neuroscience, 6(8):2240–2244, 1986.

8. Soares, A. F., Nissen, J. D., Garcia-Serrano, A. M., Nussbaum, S. S., Waagepetersen, H. S., and Duarte, J. M. Glycogen metabolism is impaired in the brain of male type 2 diabetic goto-kakizaki rats. Journal of Neuroscience Research, 97(8): 1004–1017, 2019.

9. Baker, L. D., Cross, D. J., Minoshima, S., Belongia, D., Watson, G. S., and Craft, S. Insulin resistance and alzheimer-like reductions in regional cerebral glucose metabolism for cognitively normal adults with prediabetes or early type 2 diabetes. Archives of neurology, 68(1):51–57, 2011.

10. Beeri, M. S., Goldbourt, U., Silverman, J. M., Noy, S., Schmeidler, J., Ravona-Springer, R., Sverdlick, A., and Davidson, M. Diabetes mellitus in midlife and the risk of dementia three decades later. Neurology, 63(10):1902–1907, 2004.

11. Antal, B., McMahon, L. P., Sultan, S. F., Lithen, A., Wexler, D. J., Dickerson, B., Ratai, E.-M., and Mujica-Parodi, L. R. Type 2 diabetes mellitus accelerates brain aging and cognitive decline: Complementary findings from uk biobank and meta-analyses. Elife, 11:e73138, 2022.

12. Cunnane, S. C., Courchesne-Loyer, A., St-Pierre, V., Vandenberghe, C., Pierotti, T., Fortier, M., Croteau, E., and Castellano, C.-A. Can ketones compensate for deteriorating brain glucose uptake during aging? implications for the risk and treatment of alzheimer’s disease. Annals of the New York Academy of Sciences, 1367(1):12–20, 2016.

13. Zilberter, Y. and Zilberter, M. The vicious circle of hypometabolism in neurodegenerative diseases: ways and mechanisms of metabolic correction. Journal of neuroscience research, 95(11):2217–2235, 2017.

14. Mujica-Parodi, L. R., Amgalan, A., Sultan, S. F., Antal, B., Sun, X., Skiena, S., Lithen, A., Adra, N., Ratai, E.-M., Weistuch, C., et al. Diet modulates brain network stability, a biomarker for brain aging, in young adults. Proceedings of the National Academy of Sciences, 117(11):6170–6177, 2020.

15. Weistuch, C., Mujica-Parodi, L. R., Razban, R. M., Antal, B., van Nieuwenhuizen, H., Amgalan, A., and Dill, K. A. Metabolism modulates network synchrony in the aging brain. Proceedings of the National Academy of Sciences, 118(40):e2025727118, 2021.

16. Pantano, P., Baron, J.-C., Lebrun-Grandié, P., Duquesnoy, N., Bousser, M.-G., and Comar, D. Regional cerebral blood flow and oxygen consumption in human aging. Stroke, 15(4):635–641, 1984.

17. Hasselbalch, S. G., Madsen, P. L., Hageman, L. P., Olsen, K. S., Justesen, N., Holm, S., and Paulson, O. B. Changes in cerebral blood flow and carbohydrate metabolism during acute hyperketonemia. American Journal of Physiology-Endocrinology And Metabolism, 270(5):E746–E751, 1996.

18. Logothetis, N., Pauls, J., Augath, M., Trinath, T., and Oeltermann, A. Neurophysiological investigation of the basis of the fmri signal. Nature, 412:150–157, 2001.

19. Breakspear, M., Terry, J. R., and Friston, K. J. Modulation of excitatory synaptic coupling facilitates synchronization and complex dynamics in a biophysical model of neuronal dynamics. Network: Computation in Neural Systems, 14(4):703, 2003.

20. da Silva, F. L. Eeg and meg: relevance to neuroscience. Neuron, 80(5):1112–1128, 2013.

21. Jensen, N. J., Wodschow, H. Z., Nilsson, M., and Rungby, J. Effects of ketone bodies on brain metabolism and function in neurodegenerative diseases. International journal of molecular sciences, 21(22):8767, 2020.

22. Hoyer, S. The abnormally aged brain. its blood flow and oxidative metabolism. a review—part ii. Archives of gerontology and geriatrics, 1(3):195–207, 1982.

23. Ma, W., Berg, J., and Yellen, G. Ketogenic diet metabolites reduce firing in central neurons by opening katp channels. Journal of Neuroscience, 27(14):3618–3625, 2007.

24. Chesebro, A. G., Mujica-Parodi, L. R., and Weistuch, C. Ion gradient-driven bifurcations of a multi-scale neuronal model. Chaos, Solitons & Fractals, 167:113120, 2023.

25. Baeza-Lehnert, F., Saab, A. S., Gutiérrez, R., Larenas, V., Díaz, E., Horn, M., Vargas, M., Hösli, L., Stobart, J., Hirrlinger, J., et al. Non-canonical control of neuronal energy status by the na+ pump. Cell metabolism, 29(3):668–680, 2019.

26. Sesti, F., Liu, S., and Cai, S.-Q. Oxidation of potassium channels by ros: a general mechanism of aging and neurodegeneration? Trends in cell biology, 20(1):45–51, 2010.

27. Scott, B., Leu, J., and Cinader, B. Effects of aging on neuronal electrical membrane properties. Mechanisms of ageing and development, 44(3):203–214, 1988.

28. Farajnia, S., Meijer, J. H., and Michel, S. Age-related changes in large-conductance calcium-activated potassium channels in mammalian circadian clock neurons. Neurobiology of aging, 36(6):2176–2183, 2015.

29. Power, J. M., Wu, W. W., Sametsky, E., Oh, M. M., and Disterhoft, J. F. Age-related enhancement of the slow outward calcium-activated potassium current in hippocampal ca1 pyramidal neurons in vitro. Journal of Neuroscience, 22(16): 7234–7243, 2002.

30. Larter, R., Speelman, B., and Worth, R. M. A coupled ordinary differential equation lattice model for the simulation of epileptic seizures. Chaos: An Interdisciplinary Journal of Nonlinear Science, 9(3):795–804, 1999.

31. Roberts, J. A., Gollo, L. L., Abeysuriya, R. G., Roberts, G., Mitchell, P. B., Woolrich, M. W., and Breakspear, M. Metastable brain waves. Nature communications, 10(1):1–17, 2019.

32. Byrne, Á., O’Dea, R. D., Forrester, M., Ross, J., and Coombes, S. Next-generation neural mass and field modeling. Journal of neurophysiology, 123(2):726–742, 2020.

33. Meyer, D. J., Díaz-García, C. M., Nathwani, N., Rahman, M., and Yellen, G. The na+/k+ pump dominates control of glycolysis in hippocampal dentate granule cells. Elife, 11:e81645, 2022.

34. Peirce, J. W. Psychopy—psychophysics software in python. Journal of neuroscience methods, 162(1-2):8–13, 2007.

35. Oostenveld, R. and Praamstra, P. The five percent electrode system for high-resolution eeg and erp measurements. Clinical neurophysiology, 112(4):713–719, 2001.

36. Babayan, A., Erbey, M., Kumral, D., Reinelt, J. D., Reiter, A. M., Röbbig, J., Schaare, H. L., Uhlig, M., Anwander, A., Bazin, P.-L., et al. A mind-brain-body dataset of mri, eeg, cognition, emotion, and peripheral physiology in young and old adults. Scientific data, 6(1):1–21, 2019.

37. Delorme, A. and Makeig, S. Eeglab: an open source toolbox for analysis of single-trial eeg dynamics including independent component analysis. Journal of neuroscience methods, 134(1):9–21, 2004.

38. Pion-Tonachini, L., Kreutz-Delgado, K., and Makeig, S. Iclabel: An automated electroencephalographic independent component classifier, dataset, and website. NeuroImage, 198:181–197, 2019.

39. Gramfort, A., Luessi, M., Larson, E., Engemann, D. A., Strohmeier, D., Brodbeck, C., Goj, R., Jas, M., Brooks, T., Parkkonen, L., and Hämäläinen, M. S. MEG and EEG data analysis with MNE-Python. Frontiers in Neuroscience, 7(267):1–13, 2013. doi: 10.3389/fnins.2013.00267.

40. König, T., Lehmann, D., Saito, N., Kuginuki, T., Kinoshita, T., and Koukkou, M. Decreased functional connectivity of eeg theta-frequency activity in first-episode, neuroleptic-naive patients with schizophrenia: preliminary results. Schizophrenia research, 50(1-2):55–60, 2001.

41. Michels, L., Lüchinger, R., Koenig, T., Martin, E., and Brandeis, D. Developmental changes of bold signal correlations with global human eeg power and synchronization during working memory. PLoS One, 7(7):e39447, 2012.

42. König, T., Prichep, L., Dierks, T., Hubl, D., Wahlund, L., John, E., and Jelic, V. Decreased eeg synchronization in alzheimer’s disease and mild cognitive impairment. Neurobiology of aging, 26(2):165–171, 2005.

43. Achermann, P., Rusterholz, T., Dürr, R., König, T., and Tarokh, L. Global field synchronization reveals rapid eye movement sleep as most synchronized brain state in the human eeg. Royal Society open science, 3(10):160201, 2016.

44. Nicolaou, N. and Georgiou, J. Global field synchrony during general anaesthesia. British Journal of Anaesthesia, 112(3): 529–539, 2014.

45. Endo, H., Hiroe, N., and Yamashita, O. Evaluation of resting spatio-temporal dynamics of a neural mass model using resting fmri connectivity and eeg microstates. Frontiers in computational neuroscience, 13:91, 2020.

46. Hutchison, R. M., Womelsdorf, T., Allen, E. A., Bandettini, P. A., Calhoun, V. D., Corbetta, M., Della Penna, S., Duyn, J. H., Glover, G. H., Gonzalez-Castillo, J., et al. Dynamic functional connectivity: promise, issues, and interpretations. Neuroimage, 80:360–378, 2013.

47. Seto, E., Sela, G., McIlroy, W. E., Black, S. E., Staines, W. R., Bronskill, M. J., McIntosh, A. R., and Graham, S. J. Quantifying head motion associated with motor tasks used in fmri. Neuroimage, 14(2):284–297, 2001.

48. Taher, H., Avitabile, D., and Desroches, M. Bursting in a next generation neural mass model with synaptic dynamics: a slow–fast approach. Nonlinear Dynamics, pages 1–25, 2022.

