## Supplemental Information for "Ketosis regulates K^+^ ion channels, strengthening brain-wide signaling disrupted by age"

### Supplementary Information

#### Clinical and Demographic Information of Metabolic Study Participants

| **Measure** | | |
|---|---|---|
| **Sex** | | N = 36 (18 female) |
| **Age (years)** | Average | 26.9 ± 11.2 |
| | Median | 21.5 |
| | Range | 19 - 65 |
| **Ethnicity** | White | 16 |
| | Black | 1 |
| | Asian | 15 |
| | Hispanic | 4 |
| **HbA1c (%)** | Average | 5.1 ± 0.4 |
| | Median | 5.1 |
| | Range | 4.4 - 6.1 |

**Table S1.** Clinical and Demographic Information of the Metabolic Study Participants.

**Justification of Choice of Neural Synchrony as Macroscale Biomarker for Top-Level of Multi-Scale Model**

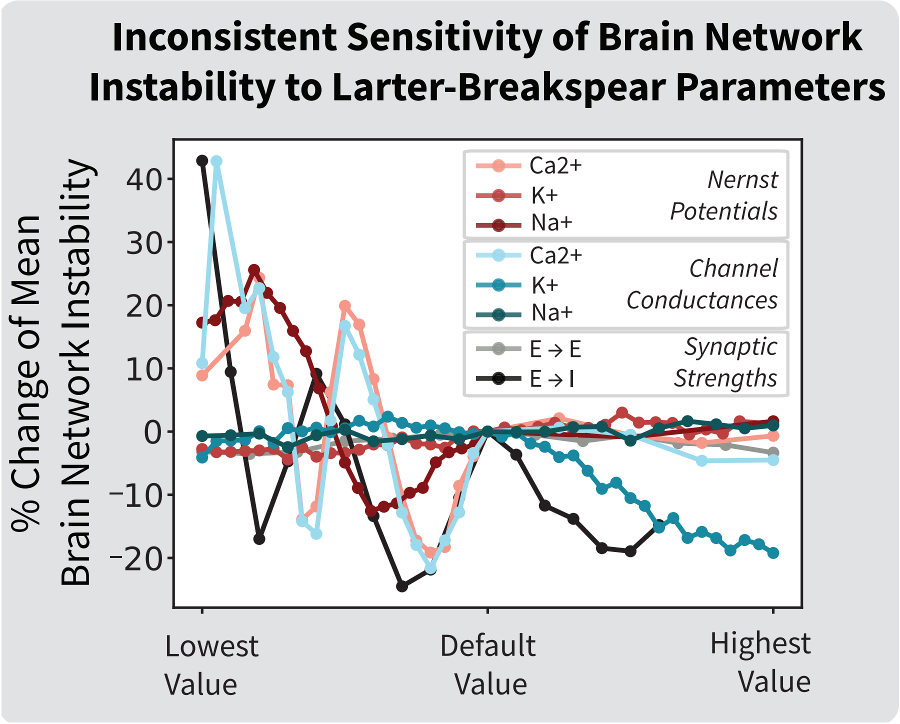


**Figure S1. Modeling brain network instability using simulated rsEEG produced by the Larter-Breakspear neural mass model shows inconsistent sensitivity of network instability to model parameter variations.**

**Effect of Window Size on Brain Network Instability Calculation**

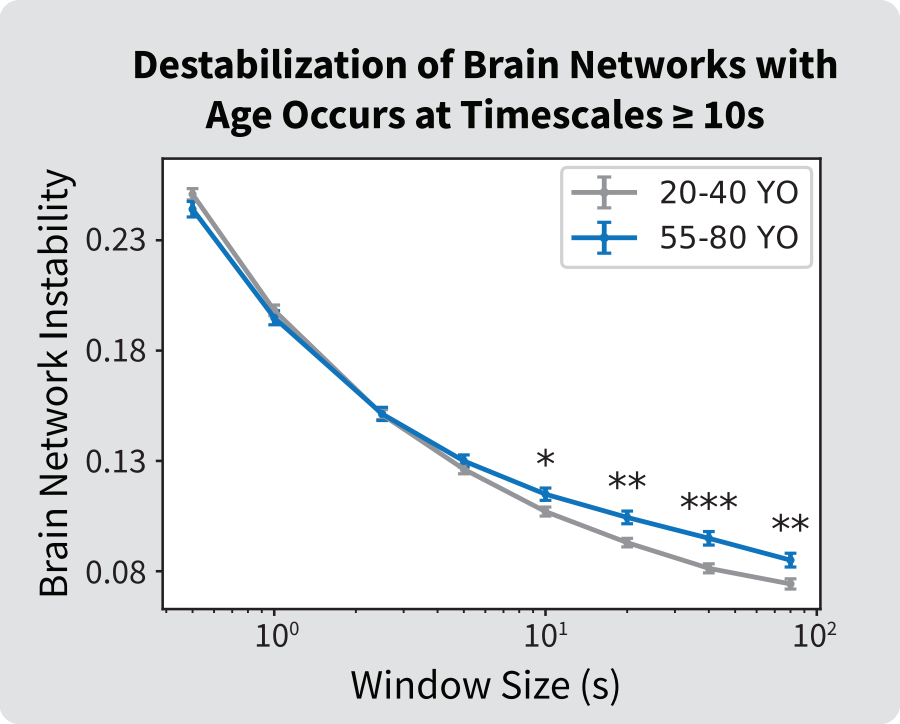


**Figure S2. High temporal resolution of EEG reveals timescale of age-related brain network destabilization.** (A) One advantage of EEG over fMRI is a more than 3,000-fold increase in temporal resolution of the measured signal. Taking advantage of this high temporal resolution, we calculated brain network instability using Leipzig LEMON's younger (N = aged 20 to 40 years) and older (aged 55 to 80 years) eyes-closed, rsEEG cohorts for a variety of window sizes ranging from 0.5 s to 80 s. We found that network instability is significantly greater in the older cohort for window sizes $\geq$ 10 s.
